# Closed-loop auditory stimulation during sleep shapes cortical responses and memory consolidation depending on spindle timing

**DOI:** 10.64898/2026.05.29.728629

**Authors:** Tao Xia, Xingyi Jin, Danni Chen, Xibo Zuo, Yuanwei Yao, Yuan Zhang, Tifei Yuan, Zhe Zhang, Xiaoli Li, Cora S.W. Lai, Xiaoqing Hu

**Affiliations:** State Key Laboratory of Cognitive Science and Mental Health, Institute of Psychology, Chinese Academy of Sciences, Beijing, China; Department of Psychology, The University of Hong Kong, Hong Kong SAR, China; Southwest University, Chongqing, China; Shanghai Key Laboratory of Psychotic Disorders, Brain Health Institute, National Center for Mental Disorders, Shanghai Mental Health Center, Shanghai Jiao Tong University School of Medicine and School of Psychology, Shanghai, 200030, China; Institute of Neuroscience, CAS Center for Excellence in Brain Science and Intelligence Technology, Chinese Academy of Sciences, Shanghai, China; School of Automation Science and Engineering, South China University of Technology, Guangzhou 510641, China; Pazhou Lab, Guangzhou 510335, China; School of Biomedical Sciences, The University of Hong Kong, Hong Kong SAR, China; HKU-Shenzhen Institute of Research and Innovation, Shenzhen, China

**Keywords:** Sleep spindles, closed-loop stimulation, auditory stimulation, memory consolidation, refractory period

## Abstract

Sleep spindles contribute to memory consolidation, and acoustic stimulation during sleep can modulate spindle activity. However, it remains unclear whether these effects depend on the timing of stimulation relative to the spindle cycle. Using closed-loop spindle detection, we delivered four variants of pink noise either during ongoing spindles or shortly after spindle offset. In Experiment 1, stimulation delivered during a spindle prolonged its duration, whereas stimulation applied shortly after spindle offset, i.e., within the refractory period, delayed the onset of the subsequent spindle. This pattern was consistent across all sound variants, indicating that the stimulation timing, rather than acoustic properties, drove the observed effects. Experiment 2 examined whether post-spindle stimulation influences memory consolidation. Relative to a no-stimulation nap, post-spindle stimulation reduced post-nap deterioration in motor sequence accuracy but did not significantly affect word-pair recall. EEG analyses revealed that stimulation elicited widespread slow oscillation activity without triggering new spindles, demonstrating that cortical processing of external input persists even during the refractory period. Notably, the motor benefit was greatest in participants with lower spindle density during the baseline adaptation nap. Together, these findings challenge the notion that the post-spindle refractory period is a functionally silent interval. Instead, this period remains receptive to external input, capable of eliciting cortical activity and supporting motor memory consolidation even in the absence of new spindle generation.

## Introduction

Sleep stabilizes newly formed memories via systems-level consolidation, a process supported by non-rapid eye movement (NREM) oscillations such as slow oscillations and sleep spindles (Born & Wilhelm, 2012; Diekelmann & Born, 2010; Rasch & Born, 2013; Stickgold, 2005). Sleep spindles are 12-16 Hz rhythmic bursts generated primarily in the thalamus, and they coordinate thalamocortical activity during NREM sleep (Astori et al., 2013; Bonjean et al., 2012; Contreras et al., 1997; Fernandez & Lüthi, 2020; Fogel & Smith, 2011; Steriade, 2003; Steriade et al., 1993). During post-learning sleep, higher spindle activity over task-related cortical regions and more spindle-rich NREM sleep have been linked to stronger memory consolidation (Barakat et al., 2011; Clemens et al., 2005; Denis et al., 2021; Morin et al., 2008; Schabus et al., 2004).

These correlational findings raise a causal question: can changing spindle activity impact memory consolidation? Closed-loop stimulation addresses this question by delivering acoustic or electrical pulses at defined points in the spindle cycle, allowing stimulation timing to be experimentally manipulated (Choi & Jun, 2022; Lustenberger et al., 2016). Previous work shows that the spindle response depends on when a stimulus arrives. Pulses near spindle onset can truncate the ongoing oscillation, whereas pulses in the waning phase can leave the detected spindle largely intact and produce a later sigma response (Jourde, Sobral, et al., 2025; Jourde, Ujevco, et al., 2025). A key question is what happens when stimulation is delivered shortly after a spindle has ended. After spindle termination, the brain enters a refractory period of approximately 1-2 s in which new spindle generation is suppressed (Antony et al., 2018; Bal & McCormick, 1996; Bazhenov et al., 2002; Fernandez & Lüthi, 2020). This suppression does not necessarily mean that the cortex is unresponsive. Motor output is reduced around spindles, as shown by decreased corticospinal excitability (Hassan et al., 2025), but auditory responses measured with time-resolved EEG/MEG are largely preserved during and after spindles, even though hemodynamic responses to tones are attenuated during spindles (Dang-Vu et al., 2011; Jourde & Coffey, 2024). Thus, stimulation during a spindle and stimulation shortly after spindle offset may probe different physiological states: one interacts with an active thalamocortical oscillation, whereas the other arrives when new spindles are suppressed but auditory cortical processing remains available. How these two timing conditions affect spindle dynamics within the same experimental framework, and whether post-spindle stimulation can influence memory without producing new spindles, remains unknown.

The behavioral consequences of spindle phase-dependent stimulation have been studied mainly when stimulation is delivered during, or close to, active spindles. Auditory and electrical stimulation during this time window can benefit memory consolidation (Baxter et al., 2023; Choi & Jun, 2022; Kasties et al., 2025; Lustenberger et al., 2016). More broadly, targeted memory reactivation and closed-loop stimulation studies show that external cues timed to endogenous sleep rhythms, especially slow oscillation up-states, can influence subsequent memory (Besedovsky et al., 2017; Esfahani et al., 2023; Geva-Sagiv et al., 2023; Hu et al., 2020; Leminen et al., 2017; Marshall et al., 2006; Ngo et al., 2013; Ong et al., 2016; Santostasi et al., 2016). Yet these approaches leave open whether stimulation remains effective immediately after a spindle has ended. This post-spindle window is theoretically important because it coincides with a refractory period in which new spindle generation is temporarily suppressed, while auditory processing may remain available at the cortical level (Antony et al., 2018; Jourde & Coffey, 2024; Jourde et al., 2025a).

In addition to stimulation timing, acoustic properties, especially the duration and sound envelope, may play an important role. Previous closed-loop studies used pink-noise pulses of 15 to 50 ms (Choi & Jun, 2022; Choi et al., 2019; Jourde et al., 2026; Jourde, Sobral, et al., 2025; Ngo et al., 2013), shorter than a single spindle cycle. These brief pulses are useful for testing immediate, phase-locked perturbations, but they cannot determine whether longer acoustic input changes spindle duration or the timing of the next spindle. The sound envelope may also shape the response. For example, white noise amplitude-modulated at 12 Hz increased slow spindle activity, whereas 15 Hz modulation increased fast spindle activity (Antony & Paller, 2017), suggesting that spindle responses may depend on how closely the acoustic envelope matches endogenous sigma rhythms. Another study used spindle-frequency rhythmic pulses, but the input lasted only about half a spindle cycle and did not sustain the ongoing oscillation (Ngo et al., 2019). It is therefore still unknown whether longer stimulation spanning multiple spindle cycles can prolong ongoing spindles or extend the post-spindle refractory period.

Here, we conducted two experiments to test how stimulation timing relative to the spindle cycle affects spindle dynamics and memory consolidation. We implemented a closed-loop spindle stimulation procedure to deliver 2 s pink-noise bursts either during an ongoing spindle, at the fifth oscillatory peak after online detection, or 250 ms after spindle offset, within the post-spindle refractory period (Antony et al., 2018). Experiment 1 tested whether acoustic properties interacted with timing by comparing four pink-noise variants. Three variants were structured at each participant’s individualized spindle frequency (sigma-band amplitude-modulated, rhythmic, and arrhythmic), and the fourth used a 30 Hz control frequency. We found that during-spindle stimulation prolonged the ongoing spindle, whereas post-spindle stimulation delayed the next spindle. This dissociation was present across all four sound types, indicating that the stimulation timing, rather than acoustic properties, drove the observed effects. Experiment 2 tested whether the post-spindle stimulation effects identified in Experiment 1 could influence memory consolidation. Because cortical responses to sound can persist during the refractory period (Jourde & Coffey, 2024), post-spindle stimulation might affect memory without generating new spindles. Experiment 2 thus used a within-participant design comparing a stimulation nap with an adaptation nap and showed that post-spindle stimulation reduced the post-nap loss of finger-tapping accuracy and elicited widespread slow-oscillation activity without generating new spindles. In contrast, post-spindle stimulation had no detectable effect on word-pair recall. The motor benefit was largest in participants with lower endogenous spindle activity, suggesting that external stimulation may be most useful when spontaneous spindle activity is relatively weak.

## Methods

### 2.1 Participants

Participants were recruited through university advertisements. Inclusion criteria were normal self-reported hearing, no habitual use of earplugs during sleep, and no history of neurological or psychiatric conditions. Participants abstained from alcohol, caffeine, tobacco, and medications on experimental days. The study was approved by the Ethics Committee of the University of Hong Kong (EA210341); all participants provided written informed consent.

#### Experiment 1

Forty-one participants were enrolled. Nineteen were excluded: 6 withdrew after a poor adaptation nap, 4 had prolonged wakefulness during the stimulation nap (>10 min after stimulation had begun), and 9 obtained insufficient NREM sleep to complete all four acoustic conditions. The final sample comprised 22 participants (21 females; age 21 ± 2 years, mean ± SD), randomly assigned to the During-spindle (n = 11) or Post-spindle (n = 11) group.

#### Experiment 2

Forty-five participants were enrolled. Eighteen were excluded: 7 withdrew after inadequate adaptation-nap sleep, 4 received fewer than the required minimum of 40 stimulations, and 7 had excessive EEG artifacts. The retained sample comprised 27 participants (21 females; age 22 ± 2 years, mean ± SD). One participant exceeded ±3 SD from the sample mean on the Day 2 − Day 1 finger-tapping accuracy difference (Figure S3) and was excluded from behavioral analyses. A second participant had a corrupted Day 1 EEG recording and was excluded from cross-session analyses but retained for Day 2 EEG analyses. Final analysis samples: N = 26 for behavioral comparisons, N = 26 for Day 2 EEG analyses, and N = 25 for Day 1 versus Day 2 comparisons and cross-session EEG–behavior correlations.

Sample size and statistical power. No a priori power calculation was performed; sample sizes were guided by recent sleep closed-loop studies (10-24 participants per group; Lustenberger et al., 2016; Ngo et al., 2019; Baxter et al., 2023; Jourde et al., 2025a; Kasties et al., 2025). Post hoc sensitivity analyses (G*Power 3.1.9.7; two-tailed α = .05, 1 - β = .80) indicated that Experiment 1 (n = 11 per group, independent-samples t-test) had 80% power to detect between-group effects of d ≥ 1.25. In Experiment 2 (paired t-test), 80% power was achieved for within-participant effects of dz ≥ 0.57 for behavioral comparisons (N = 26) and dz ≥ 0.58 for cross-session EEG comparisons (N = 25), and for EEG-behavior correlations of |r| ≥ 0.52 (N = 25). Smaller effects may not have been detected.

### 2.2 Experimental Protocols

Both experiments used a two-day nap protocol: an adaptation nap on Day 1 (no stimulation) followed by a stimulation nap on Day 2, each lasting 90 minutes and beginning at approximately 12:30 p.m. (Figure 1A,B). The adaptation nap was used to calibrate individualized spindle-detection parameters (peak sigma frequency and amplitude threshold; see Section 2.4) and to acclimate participants to the laboratory. Trained experimenters performed real-time sleep staging from Cz, F3/4, PO3/4, EOG, and EMG, using visual AASM criteria (Berry et al., 2012; Iber et al., 2007). Stimulation began after 5 minutes of continuous NREM sleep. Participants were awakened after 90 minutes of recording, or earlier if they remained awake for more than 10 minutes after stimulation had begun. This criterion limited prolonged wakefulness during the nap; it should not be interpreted as evidence that stimulation caused awakening.

**Figure 1.**
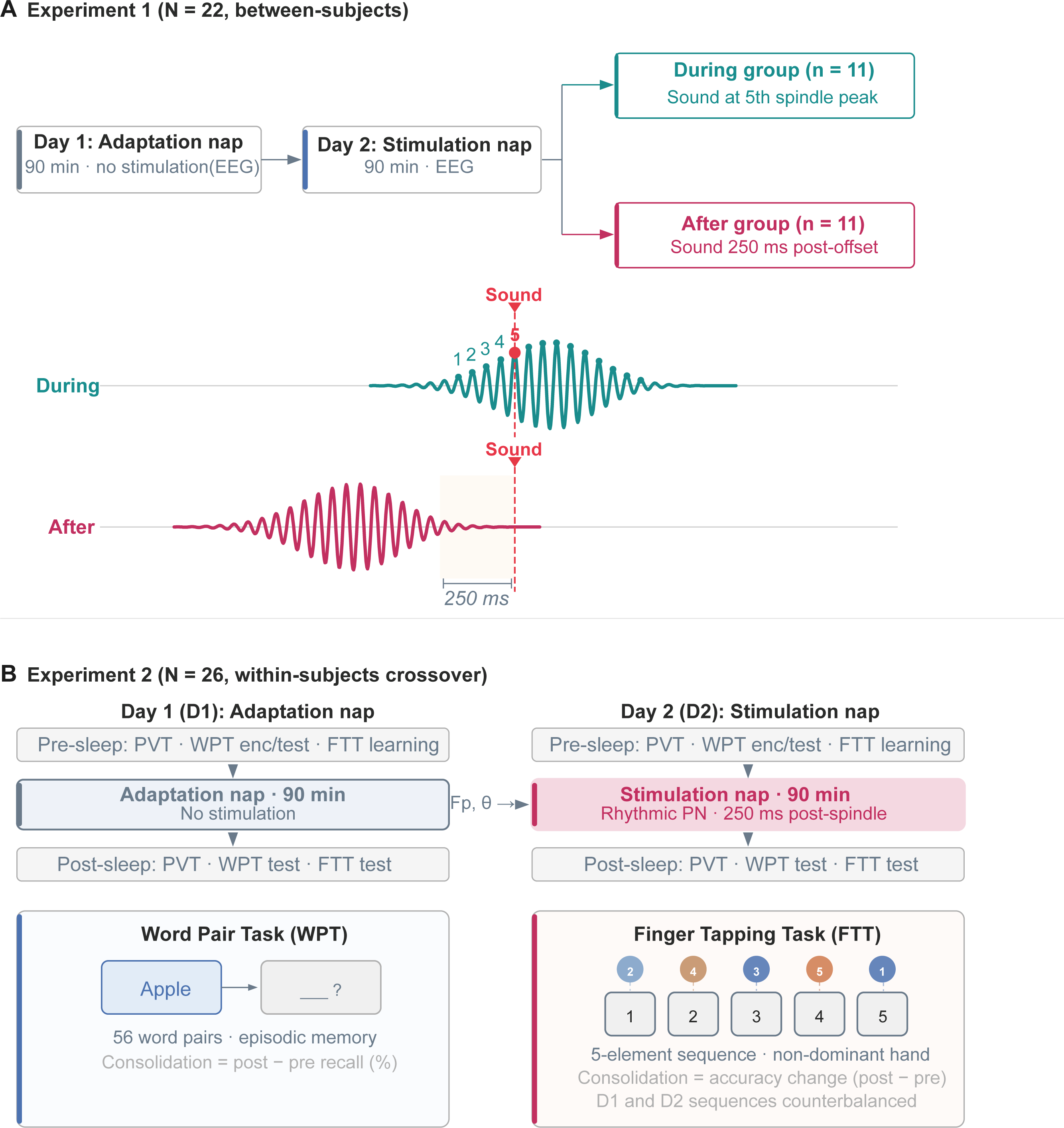
Experimental design. (A) Experiment 1 (N = 22, between-subjects). Top: participants completed an adaptation nap (Day 1) followed by a stimulation nap (Day 2) and were assigned to the During-spindle group (n = 11, stimuli delivered at the 5th spindle peak) or the Post-spindle group (n = 11, stimuli delivered 250 ms after spindle offset). Bottom: schematic spindle waveforms illustrating the two timing conditions. Waveform shape reflects the average spindle-locked waveform (∼13 Hz). (B) Experiment 2 (N = 27 enrolled, N = 26 analyzed; within-subjects). Top: all participants completed an adaptation nap (Day 1, no stimulation) followed by a stimulation nap (Day 2, post-spindle rhythmic pink noise at the individualized sigma frequency). Pre- and post-nap assessments included the PVT, WPT, and FTT. Bottom: task schematics. WPT: 56 word pairs, cued recall. FTT: 5-element motor sequence, non-dominant hand; Day 1 and Day 2 sequences counterbalanced.

#### Experiment 1

Stimulation timing and spindle dynamics. Experiment 1 tested how stimulation timing relative to the spindle cycle shapes spindle dynamics, using a between-subjects design (Figure 1A). Participants were randomly assigned to the During-spindle group (n = 11), which received auditory stimuli at the fifth peak of detected spindles to maximize interaction with the ongoing oscillation (Lustenberger et al., 2016), or the Post-spindle group (n = 11), which received stimuli 250 ms after spindle offset (Antony et al., 2018). The 250 ms delay ensured that the stimulus arrived after the spindle had ended, providing a clean separation from the during-spindle condition. Four pink-noise variants were delivered in a pseudorandomized block design to test whether acoustic properties interacted with stimulation timing. Each block contained 10 stimuli of a single sound type, and block order was pseudorandomized within each participant, with the constraint that all four sound types occurred once before any type was repeated. The four variants were sigma-band amplitude-modulated, 30 Hz amplitude-modulated (control frequency), rhythmic, and arrhythmic pink noise (acoustic details in Section 2.4). Blocks were separated by 30 s rest periods, and stimuli within a block were delivered with a minimum 5 s inter-stimulus interval.

#### Experiment 2

Post-spindle stimulation and tests for memory consolidation. Experiment 2 tested whether post-spindle stimulation supports memory consolidation, using a within-participant design that compared the Day 2 stimulation nap with the Day 1 adaptation nap (Figure 1B). A fixed order was necessary because the individualized spindle-detection parameters used on Day 2 had to be derived from the Day 1 adaptation nap. Stimulation used only the rhythmic pink-noise variant, delivered at the participant’s individualized sigma-peak frequency 250 ms after spindle offset, with the same block structure as Experiment 1. Pre- and post-nap assessments included the Psychomotor Vigilance Task (PVT), the word-pair task (WPT), and the finger-tapping task (FTT; see Section 2.3). To control for practice effects on the FTT, each participant learned a different five-element motor sequence on Day 1 and Day 2, with sequence assignments counterbalanced across participants.

### 2.3 Memory Tasks

Two tasks were used to evaluate sleep-dependent memory consolidation: the WPT for episodic memory and the FTT for procedural motor memory.

#### Word-pair task (WPT)

Participants learned 56 semantically unrelated Chinese word pairs presented visually on a computer screen. Each pair appeared for 5 s with a 2 s inter-pair interval, and the full list was presented twice. Half of the word pairs were encoded before the adaptation nap and half before the stimulation nap, with the assignment of pair sets to naps counterbalanced across participants. Immediate and post-nap recall used the same cued-recall procedure: one word of each pair appeared on screen and participants had 4 s to verbally recall the paired associate. Post-nap recall began immediately upon waking. The primary outcome was the percentage-point change in recall accuracy from immediate to post-nap recall.

#### Finger-tapping task (FTT)

Participants learned a five-element sequence (e.g., 4-1-3-2-4) by typing it repeatedly on a keyboard with their non-dominant hand (Karni et al., 1994; Walker et al., 2002; Walker et al., 2003). Pre-nap training consisted of twelve 30 s tapping blocks separated by 30 s rest periods; post-nap testing used the same structure. Performance was quantified as accuracy (correct sequences / total sequences) and reaction time per correct sequence. To minimize sequence-specific learning transfer between days in Experiment 2, two equivalent sequences were counterbalanced across participants and sessions.

### 2.4 Closed-Loop Auditory Stimulation and EEG Recording

#### EEG acquisition

Sleep EEG was recorded with a 64-channel system (eego, ANT Neuro, The Netherlands) following the international 10–20 system, with simultaneous electrooculography (EOG) and electromyography (EMG). Electrode impedances were maintained below 20 kΩ. For closed-loop processing, signals were acquired via OpenViBE (v3.4.0) at 500 Hz and downsampled online to 250 Hz. Cz served as the detection channel and CPz as the online reference.

Real-time spindle detection. The Cz signal was broadband-filtered (0.1-40 Hz) and then band-pass filtered around the participant’s individualized peak sigma frequency (peak frequency ± 1.5 Hz; 2nd-order Butterworth). The sigma envelope was constructed by interpolating consecutive positive peaks of the filtered signal rather than by using the Hilbert transform, which reduced total system latency to <50 ms. A spindle was detected when the envelope exceeded an individualized amplitude threshold for 0.5-3 s, consistent with standard criteria (Berry et al., 2012).

#### Parameter calibration

Detection parameters were individualized using the adaptation nap recording (Carrier et al., 2001; Chen et al., 2025; Lacourse et al., 2019; Purcell et al., 2017; Warby et al., 2014). The peak sigma frequency was identified as the dominant frequency in the 12-16 Hz range across N2 and N3 sleep (Figure S5). The amplitude threshold was set as Threshold = Baseline × Multiplier, where Baseline was the mean amplitude of the NREM sigma envelope and Multiplier was selected from a grid (1.0 to 2.0, step 0.1) to maximize the F1-score against the offline-detected spindle set (offline detection algorithm described in Section 2.5).

#### Auditory stimuli

Stimuli were generated in MATLAB R2024a and delivered through a loudspeaker positioned approximately 2 m from the participant’s head. Continuous pink background noise (∼37 dB SPL, measured at the head position) was played throughout each nap to mask environmental sounds. Each stimulus was a 2 s burst of pink noise (approximately matching typical spindle duration), superimposed on this background at approximately 1.5 dB above the background level. Experiment 1 used four pink-noise variants that shared the same carrier but differed in amplitude modulation: (i) sigma-band amplitude-modulated, with the carrier modulated at the participant’s individualized sigma-peak frequency (12–16 Hz, derived from Day 1 EEG); (ii) 30 Hz amplitude-modulated, modulated at a fixed supra-sigma control frequency; (iii) rhythmic, with energy pulsed at the participant’s individualized sigma-peak frequency; and (iv) arrhythmic, with modulation timing jittered to remove rhythmic structure. Experiment 2 used only the rhythmic variant.

### 2.5 Offline Data Processing and Analysis

#### EEG preprocessing

Offline analysis was performed in MNE-Python (Python 3.12, MNE 1.8.0). EEG data were re-referenced to the average of the bilateral mastoids (M1, M2), notch-filtered at 50 Hz (2nd-order Butterworth IIR) to remove power-line noise, and band-pass filtered between 0.1 and 40 Hz with a 2nd-order Butterworth filter applied in a zero-phase forward-backward manner.

#### Sleep staging

Sleep stages were initially scored using the YASA library (v0.6.0; Vallat & Walker, 2021) per AASM guidelines (Berry et al., 2012), then manually verified by two sleep researchers blinded to stimulation events, using channels Fp1/2, F3/4, C3/4, Cz, P3/4, O1/2, EMG, and EOG. Disagreements were resolved by discussion.

#### Spindle detection

Spindles were detected offline using a threshold-based algorithm (Helfrich et al., 2018) applied to the Cz channel (re-referenced to M1, M2). The signal was band-pass filtered to the participant’s individualized sigma peak (peak frequency ± 1.5 Hz; 2nd-order Butterworth), and the analytic amplitude envelope was computed via Hilbert transform and smoothed with a 200 ms moving-average window. Spindles were identified as contiguous above-threshold regions, with the threshold set at the 75th percentile of the smoothed envelope amplitude across all N2/N3 samples (Helfrich et al., 2018), and duration constrained to 0.5–3.0 s. The spindle peak was defined as the sample with maximum smoothed envelope amplitude within each event; events with absolute peak amplitude >150 µV were excluded as artifacts. This offline algorithm approximates the real-time pipeline (Section 2.4) but operates on the full continuous recording with non-causal filtering. All spindle-level analyses (duration, post-stimulation latency, EEG–behavior correlations) used this offline detection; the real-time algorithm served only to time stimulus delivery during the nap. To assess robustness to detection errors, we conducted a sensitivity analysis progressively restricting analyses to the highest-amplitude spindles (Results 3.1, Table S3).

#### Post-stimulation spindle latency

We used post-spindle latency to quantify when spindle activity returned after the target spindle. For each stimulation-locked epoch, latency was measured from the offset of the target spindle to the onset of the first subsequent spindle detected within the following 3 s. This 3 s search window was chosen to cover the expected post-spindle refractory period of approximately 1–2 s while allowing for modest inter-individual variability (Antony et al., 2018; Bal & McCormick, 1996).

Epochs without a subsequent spindle within this 3 s window did not yield a latency value and were excluded from latency analyses. This occurred in 77.3% of target epochs in the During-spindle group and 66.6% in the Post-spindle group, consistent with the fact that spontaneous inter-spindle intervals in NREM sleep are often longer than the refractory period itself. Importantly, the same 3 s search window was applied to non-target external spindles used as the within-subject baseline, so target and baseline events were censored in the same way.

The closed-loop system enforced a minimum 5.0 s interval between stimulation onsets. This restriction affected only the delivery of subsequent stimuli; it did not prevent offline detection of spindles occurring after the target spindle. Offline spindle detection resumed immediately after stimulus offset and did not include a blanking window. Thus, the latency metric was not artificially lengthened by the stimulation-spacing rule.

Within-subject baselines for control analyses. We used different within-subject baselines for the duration and latency analyses because the two measures ask different questions. For spindle duration, the relevant comparison was whether a stimulated target spindle was longer than similar unstimulated spindles from the same nap and sleep stage. We therefore used non-target spindles from missed epochs: stimulation-locked epochs that contained an offline-detected spindle but did not meet the target-selection rule; for example, epochs in which the spindle offset fell outside the ±250 ms target window. For post-spindle latency, the relevant comparison was when the next spindle occurred after a target event. Because this measure depends on the local spacing of spindles, we used non-target spindles from NREM segments outside the ±3 s stimulation-locked window, matched by participant and sleep stage, and applied the same 3 s search window. Thus, duration and latency were each compared with an unstimulated baseline matched to the structure of that measure.

#### Stimulation accuracy

Stimulation accuracy was quantified offline against the spindle set described above. For During-spindle stimulation, accuracy was the proportion of stimuli that fell within the onset-to-offset window of an offline-detected spindle. For post-spindle stimulation, accuracy was the proportion of stimuli that occurred 250 ± 250 ms after the offset of an offline-detected spindle. The during-spindle and post-spindle real-time detectors used different criteria. During-spindle detection relied on amplitude threshold alone, enabling rapid stimulus delivery during the ongoing oscillation. Post-spindle detection additionally required that the spindle had sustained above-threshold activity for a minimum duration, providing more reliable confirmation of spindle offset. Accuracy values across conditions and sleep stages are reported in Results 3.1.

### 2.6 Statistical Analysis

Statistical analyses were performed in Python (SciPy v1.13.1, pingouin v0.5, statsmodels v0.14) with a two-tailed significance threshold of α = .05. Before inferential testing, spindle measures were averaged within each participant so that analyses used subject-level means rather than repeated spindle-level observations. Outliers were defined by a ±3 SD criterion at two levels. At the trial level, FTT reaction times more than 3 SD from the participant’s block-level mean were excluded before averaging. At the participant level, Day 2 - Day 1 difference scores exceeding ±3 SD from the sample mean were excluded (n = 1; see Section 2.1).

Group and within-subject comparisons. Between-group comparisons (During-spindle vs. Post-spindle in Experiment 1) used Welch’s t-test to allow for unequal variances and group sizes. Within-subject comparisons (Day 1 vs. Day 2 in Experiment 2; target vs. non-target spindles within group) used paired t-tests. A mixed-design ANOVA (between factor: group; within factor: sound type) assessed effects of stimulus type on spindle duration in Experiment 1. Effect sizes are reported to three decimal places: Cohen’s d (pooled SD) for independent samples, Cohen’s dz (SD of within-subject differences) for paired samples, partial eta-squared (η²p) for ANOVA, and Pearson’s r for correlations. FTT accuracy, FTT reaction time, and WPT recall accuracy were treated as separate planned behavioral outcomes. We therefore report uncorrected p-values for these planned comparisons. ANOVA post-hoc pairwise comparisons, when reported, were controlled using the Benjamini-Hochberg FDR procedure.

#### Bayesian analysis

For null findings on primary outcomes, Bayes factors (BF₁₀) were computed in pingouin (v0.5), comparing the observed data under the alternative versus the null hypothesis. BF₁₀ < 1/3 was interpreted as moderate evidence favoring the null, BF₁₀ > 3 as moderate evidence favoring the alternative, and 1/3 ≤ BF₁₀ ≤ 3 as inconclusive (Lee & Wagenmakers, 2014).

#### Time-frequency cluster-based permutation tests

Stimulation-locked time-frequency representations (sigma and slow-oscillation bands) were tested against zero using cluster-based permutation, implemented in MNE-Python with 5,000 permutations of the sign-flipping null. Adjacent time–frequency samples exceeding a cluster-forming threshold (uncorrected p < .05) were grouped, and the empirical null was defined by the maximum summed cluster statistic across permutations. Clusters with permutation p ≤ .05 were considered significant. Channel-level FDR correction (q < .05) was applied to topographic SO power maps (Results 3.4).

#### EEG-behavior relationships

EEG-behavior correlations used Pearson’s r on subject-level data. Statistical significance was assessed with permutation testing (5,000 permutations of the behavioral variable), which generated an empirical null distribution for the observed r; permutation p-values are reported alongside Pearson’s r. We used three analyses to examine how spindle density related to motor memory. First, session-specific Pearson correlations tested the natural spindle-memory relationship separately on Day 1 (without stimulation) and Day 2 (with stimulation). Second, a linear mixed model with a density × session interaction (FTT change ∼ spindle density × session + (1 | subject)) tested whether this relationship changed between the adaptation and stimulation sessions. A negative interaction would indicate that motor consolidation depended less on endogenous spindle density during the stimulation session. Spindle density was calculated as the number of detected spindles divided by total NREM duration across the full nap, separately for Day 1 and Day 2. Third, a between-session correlation tested whether adaptation-nap spindle density predicted the stimulation benefit, defined as the Day 2 - Day 1 FTT accuracy difference.

## Results

### 3.1. During-spindle and post-spindle stimulation differentially modulated spindle activity

If during-spindle and post-spindle stimulation affect different neural states, they should change different aspects of spindle activity. We tested this with two measures from stimulation-locked epochs (±3 s around each stimulation event): target spindle duration, which tests whether the stimulation extends the ongoing oscillation, and post-spindle latency, which measures the time from target-spindle offset to the next detected spindle.

The closed-loop system delivered stimuli at the intended times in both groups (Figure 2A): spindle detection probability peaked at stimulation onset for the During-spindle group and had fallen to near zero by stimulation onset for the Post-spindle group. Quantified against the offline-detected spindle set, both groups achieved >80% targeting accuracy across all NREM stages. Accuracy was higher in the During-spindle group than in the Post-spindle group (89.8 ± 5.79% vs. 82.3 ± 6.30%, M ± SD; Welch t(19.9) = 2.91, p = .009).

**Figure 2.**
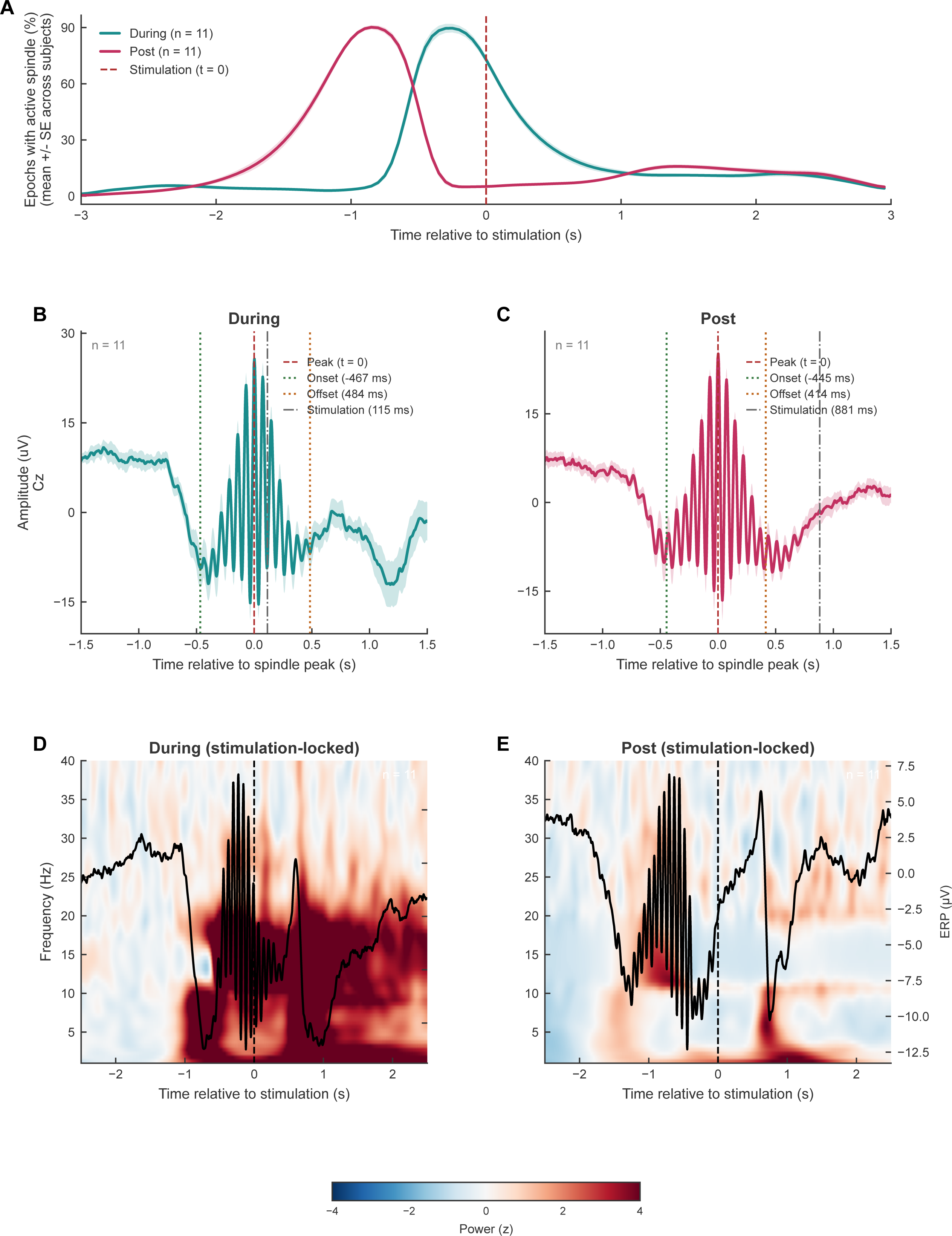
Stimulation-locked validation: spindle detection, event-related potentials, and time-frequency representations. (A) Spindle detection probability relative to stimulation onset (t = 0) for the During-spindle group (teal) and Post-spindle group (rose). Shading indicates SEM. In the During-spindle group, probability peaks at t = 0; in the Post-spindle group, probability has returned to near zero before stimulation, confirming that the target spindle ended before stimulation. (B) Grand-average broadband ERP aligned to the sigma-band envelope peak for the During-spindle group. (C) Same as (B) for the Post-spindle group. Both groups show characteristic spindle waxing-and-waning morphology. Shading indicates SEM. n = 11 per group. (D) Stimulation-locked time-frequency representation for the During-spindle group (Cz channel; Morlet wavelet, 1-40 Hz, z-scored against −2.5 to −1.0 s baseline). Black line: grand-average broadband ERP overlay. (E) Same as (D) for the Post-spindle group. Sigma-band power is elevated at stimulation onset in the During-spindle group but has returned to baseline in the Post-spindle group, confirming distinct neural states at the time of stimulation.

Spindle-locked ERPs showed the characteristic waxing-and-waning morphology of well-formed spindles in both groups (Figure 2B–C), and stimulation-locked time-frequency representations confirmed two distinct neural states at the moment of stimulation (Figure 2D–E): sigma power (12–16 Hz) was elevated at stimulation onset in the During-spindle group but had returned to baseline before stimulation in the Post-spindle group.

After verifying targeting precision, we next asked whether During-spindle stimulation extended the ongoing oscillation. Target spindles were 77 ms longer in the During-spindle group (M = 0.952 s, SD = 0.050) than in the Post-spindle group (M = 0.876 s, SD = 0.064; Welch t(18.9) = 3.14, p = .005, d = 1.338, 95% CI [0.413, 2.262]; Figure 3B-C). This between-group difference could partly reflect selection bias: stimulation in the During-spindle group was delivered at the 5th oscillatory peak (∼580 ms into the spindle on average), so shorter spindles that ended before the 5th peak would never be stimulated and would not be included in the target set.

**Figure 3.**
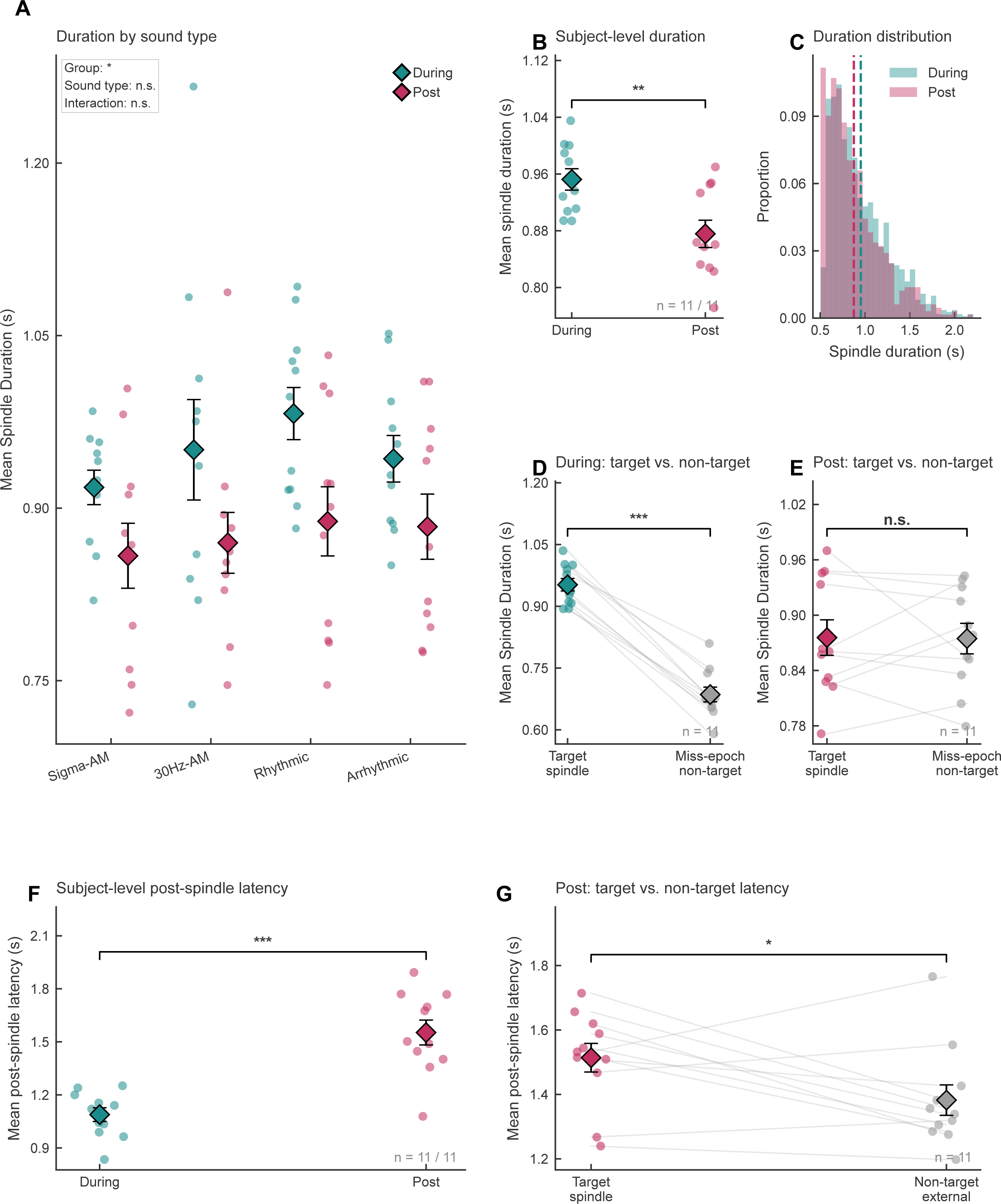
Timing-dependent spindle responses (Experiment 1). (A) Target spindle duration by sound type and group (During, teal; Post, rose). Group main effect: F(1,20) = 7.86, p = .011; no sound type effect or interaction. Individual dots represent subject means; diamonds indicate group mean ± SEM. (B) Target spindle duration: During-spindle vs. Post-spindle group. Violin plots show subject-level distributions; diamonds indicate mean ± SEM. (C) Epoch-level duration distributions for both groups. (D) During group: target spindle duration vs. non-target spindle duration in missed epochs (paired comparison). Lines connect within-subject pairs. (E) Same as (D) for the Post-spindle group. (F) Post-stimulation spindle latency: During-spindle vs. Post-spindle group. Violin plots show subject-level data. *p < .05, **p < .01, ***p < .001 (uncorrected). n = 11 per group. Error bars and diamonds indicate mean ± SEM. (G) Post-spindle group: target post-spindle latency vs. non-target external-spindle latency, matched within participant and sleep stage using the same 3 s analysis window. Lines connect within-subject pairs; diamonds indicate mean ± SEM.

To address this issue, we compared target spindles with non-target spindles from unstimulated epochs in the same nap and sleep stage, providing a within-participant baseline unaffected by acoustic input. In the During-spindle group, target spindles were still 267 ms longer than non-target spindles (0.952 s vs. 0.686 s; paired t(10) = 16.22, p < .001, dz = 4.889). In the Post-spindle group, target and non-target durations were almost identical (0.876 vs. 0.875 s; t(10) = 0.06, p = .950, dz = 0.019). Thus, this pattern cannot be explained by target selection alone. These results indicate that for both within- and between-group comparisons, during-spindle stimulation prolonged the duration of spindles beyond the expected target-selection effect.

We next asked whether post-spindle stimulation changed the timing of the next spindle. Post-spindle latency was 464 ms longer in the Post-spindle group (M = 1.553 s, SD = 0.234) than in the During-spindle group (M = 1.089 s, SD = 0.128; Welch t(15.5) = −5.76, p < .001, d = 2.46, 95% CI [1.35, 3.57]; Figure 3F). To isolate the within-subject effect, we then compared post-spindle target events with non-target spindles from external NREM segments outside the stimulation-locked epochs, matched by participant and sleep stage, using the same 3 s search window. The latency after target spindles was 132 ms longer than after non-target external spindles (paired t(10) = 2.26, p = .048, dz = 0.68; Figure 3G), indicating that post-spindle stimulation delayed the next spindle even on a within-subject basis.

We ran three control analyses to test whether factors other than stimulation timing could explain these effects. First, the effect was not specific to acoustic properties. A Sound Type × Group mixed ANOVA on target spindle duration showed no main effect of sound type (F(3,60) = 1.26, p = .297, η²p = 0.059) and no Sound Type × Group interaction (F(3,60) = 0.25, p = .864, η²p = 0.012), while the Group main effect remained significant (F(1,20) = 7.86, p = .011, η²p = 0.282; Figure 3A). Second, the duration difference could not be explained by stronger target spindles in one group. Target spindle peak amplitude did not differ between the During-spindle and Post-spindle groups (Welch t(16.7) = −0.25, p = .806, d = 0.106; Figure S1), arguing against a general increase in spindle strength. Third, the effects were not driven by low-amplitude detections near the threshold. When analyses were restricted progressively to higher-amplitude spindles, both the duration and latency effects remained significant through the top 60% amplitude threshold (duration p ≤ .034, latency p ≤ .038). At the more stringent top 40% threshold, latency remained significant (p = .006), whereas the duration effect was reduced to a non-significant trend (p = .089; Table S3).

The two groups were also matched on stimulation exposure and sleep architecture. The number of cues did not differ between groups (During: 135 ± 57; Post: 126 ± 57; t(20) = 0.38, p = .710), nor did the inter-cue interval (During: 24.7 ± 10.4 s; Post: 31.3 ± 15.6 s; t(17.4) = −1.18, p = .254). Spindle density, duration, and amplitude did not differ between the adaptation and stimulation naps (all p values > .10, N = 22; Figure S2). Sleep stage proportions were comparable (all pFDR ≥ .184; Table S1). The timing-dependent effects on spindle activity therefore reflect when stimulation arrived relative to the spindle cycle, not differences in sound content, number of stimulations, sleep quality, or detection error.

In sum, Experiment 1 revealed that stimulation timing produced two distinct effects on spindle activity: during-spindle delivery prolonged the ongoing oscillation, whereas post-spindle delivery delayed the onset of the next spindle.

### 3.2. Post-spindle stimulation preserves motor sequence accuracy

Experiment 1 showed that during-spindle and post-spindle stimulation change spindle activity in different ways. Prior work has linked stimulation during or near active spindles to memory benefits (Choi & Jun, 2022; Lustenberger et al., 2016), but whether post-spindle stimulation can influence memory consolidation remains unclear. Experiment 2 addressed this question using the word-pair task (WPT) for declarative memory and the finger-tapping task (FTT) for motor memory, comparing a stimulation nap (Day 2, post-spindle rhythmic pink noise at individualized spindle frequency) with a baseline adaptation nap (Day 1, no stimulation) in 26 participants using a within-subjects design (Figure 4).

**Figure 4.**
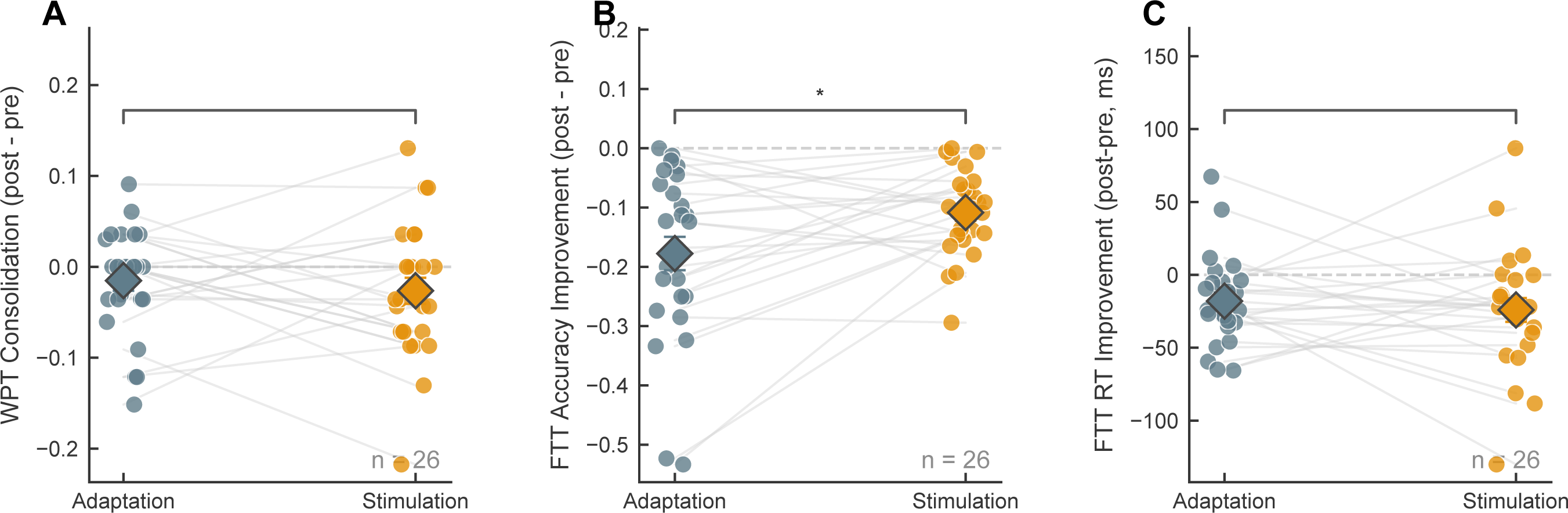
Behavioral outcomes (Experiment 2, Day 1 adaptation vs. Day 2 stimulation). (A) WPT recall change (post-nap minus pre-nap accuracy). (B) FTT accuracy change (post-nap minus pre-nap). (C) FTT reaction time change (post-nap minus pre-nap, ms). Paired violin plots show individual participant data, with lines connecting Day 1-Day 2 pairs. Diamonds indicate mean ± SEM. Each measure was analyzed as a planned behavioral outcome; uncorrected p-values are reported. Teal = Day 1 (adaptation); rose = Day 2 (stimulation). *p < .05, **p < .01. n = 26.

We compared the pre-to-post-nap change in memory accuracy between stimulation and adaptation naps as the primary behavioral outcome. For motor memory, post-spindle stimulation reduced the loss of motor sequence accuracy: both naps were followed by a decline in accuracy, but the decline was smaller after the stimulation nap (Day 1: M = −0.178, SD = 0.143; Day 2: M = −0.108, SD = 0.072; paired t(25) = −2.57, p = .016, dz = 0.505, mean difference 95% CI [0.014, 0.125]; N = 26; Figure 4B). FTT reaction time did not differ between sessions (t(25) = 0.65, p = .519, dz = −0.128, mean difference 95% CI [-25.175, 13.033] ms). For declarative memory, word-pair recall also did not differ between sessions (t(25) = 0.69, p = .496, dz = −0.136, mean difference 95% CI [-0.045, 0.022]; BF10 = 0.258, moderate evidence for the null).

We performed control analyses to test whether the Day 2 motor benefit could be explained by order-related factors in the fixed-order design. First, pre-nap FTT accuracy did not differ between Day 1 (M = 0.990, SD = 0.010) and Day 2 (M = 0.980, SD = 0.034; t(25) = 1.48, p = .151, dz = −0.291), arguing against a practice advantage on Day 2. Second, we tested whether the Day 2-Day 1 FTT difference was related to alertness, sleep duration, or baseline task performance. Pre-nap PVT reaction time, Day 2 NREM sleep duration, and Day 2 pre-nap FTT accuracy were each unrelated to the Day 2-Day 1 FTT difference (all |r| < 0.25, all p > .22). We then entered these three covariates in an adjusted regression model. The adjusted intercept remained positive and significant (b = 0.069, 95% CI [0.013, 0.125]; t(22) = 2.58, p = .017), indicating that the Day 2 preservation of motor sequence accuracy was not attributable to alertness, sleep duration, or practice carry-over from Day 1.

### 3.3. Stimulation did not alter global sleep or spindle activity

The Day 2 behavioral benefit could reflect a general difference in sleep or spontaneous spindle activity between sessions rather than a specific effect of post-spindle stimulation. If so, sleep architecture or global spindle properties should differ between Day 1 and Day 2.

In Experiment 2, continuous spindle detection applied identically to Day 1 and Day 2 revealed no differences in spindle density (Day 1: 7.19 sp/min; Day 2: 7.94 sp/min; t(24) = 1.46, p = .158, dz = 0.292), mean amplitude (t(24) = −1.10, p = .284, dz = −0.219), or mean spindle duration (Day 1: 1.039 s; Day 2: 1.024 s; t(24) = −0.66, p = .513, dz = −0.133). Within-subject variability in duration and amplitude also did not differ (duration: t(24) = 0.76, p = .457, dz = 0.151; amplitude: t(24) = 0.91, p = .369, dz = 0.183; Figure S4, N = 25). Sleep stage proportions did not differ between sessions (all FDR-corrected p values ≥ .070; Table S2). Thus, the behavioral benefit cannot be attributed to global changes in sleep architecture or spindle activity.

### 3.4. Post-spindle stimulation elicits widespread cortical activity during the spindle refractory period

After ruling out global sleep and spindle changes between sessions, we examined neural activity in the post-spindle stimulation window. Post-spindle stimulation accurately targeted the refractory period (Figure 5A-B). Sigma power was at baseline around stimulation onset, confirming low spindle activity at the time of delivery.

**Figure 5.**
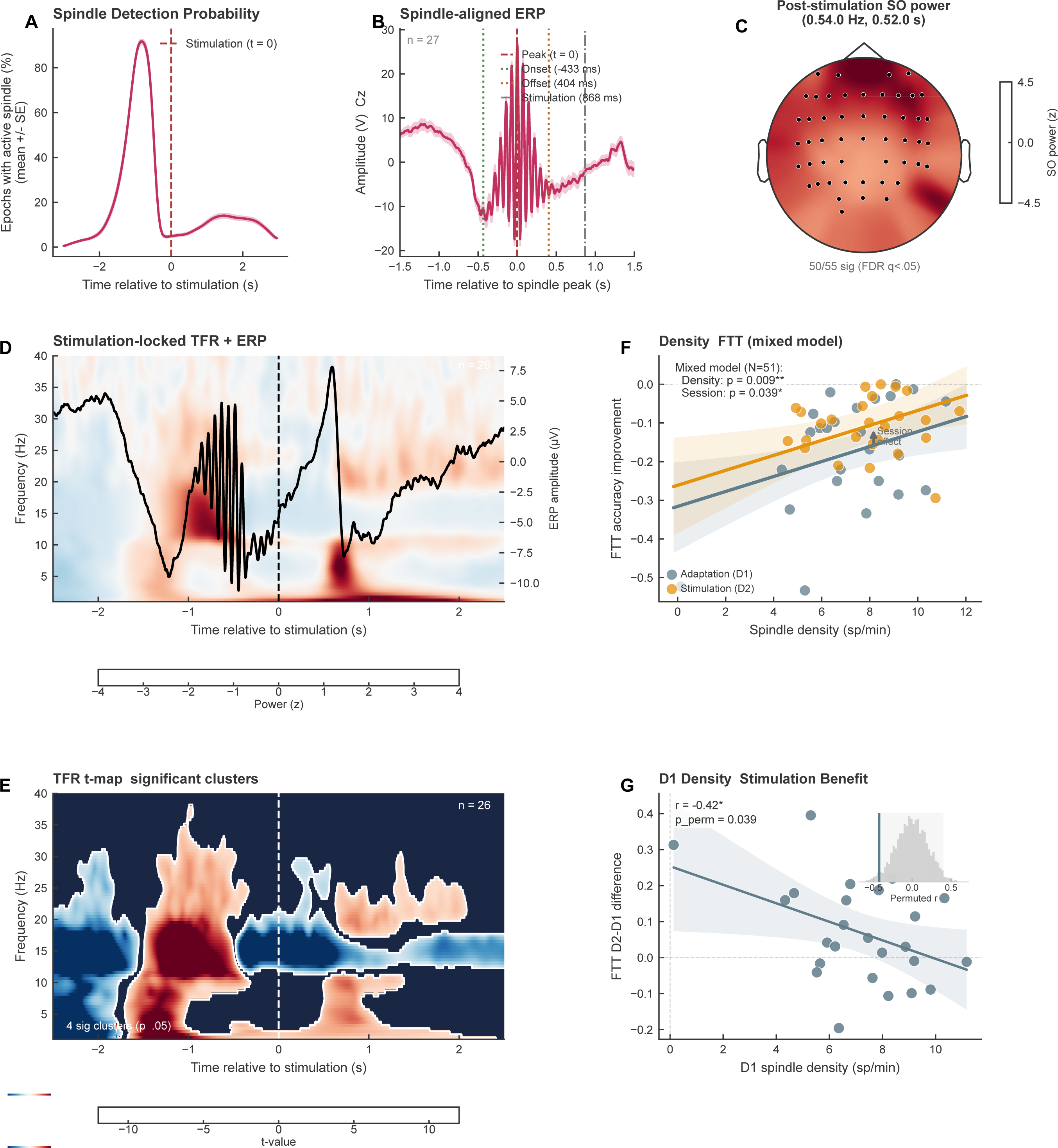
Stimulation-locked EEG responses and spindle density-motor memory relationships (Experiment 2). (A) Spindle detection probability relative to stimulation onset for Day 2 stimulation sessions. Shading indicates SEM. (B) Grand-average broadband ERP aligned to sigma-band peak (Day 2 sessions, Cz channel). Shading indicates SEM. (C) Post-stimulation slow-oscillation power (0.5-4 Hz, 0.5-2.0 s) topographic distribution. 50 of 55 channels reached significance after FDR correction (q < .05). (D) Stimulation-locked time-frequency representation (Cz channel; Morlet wavelet, 1-40 Hz, z-scored against −2.5 to −1.0 s baseline) with grand-average broadband ERP overlay (black line). Sigma power was at baseline at stimulation onset; a slow-oscillation power increase emerged after stimulation. (E) TFR t-map showing all significant clusters from the two-sided cluster-based permutation test against zero (5,000 permutations). Warm colors indicate positive t-values and cool colors indicate negative t-values. (F) Adaptation-session spindle density vs. FTT accuracy change (Day 1). Solid line shows linear regression with 95% confidence band (r = 0.508, permutation p = .010, N = 25). (G) Adaptation-session spindle density vs. stimulation benefit (Day 2-Day 1 FTT accuracy change), showing that participants with higher baseline spindle density showed smaller additional gains from stimulation (r = −0.425, permutation p = .039, N = 25). Solid lines: linear regression; shaded bands: 95% confidence intervals. Insets: permutation null distributions (5,000 iterations) with observed r indicated by vertical line. An amplitude filter (≤150 µV) was applied to all spindle data.

Even without an ongoing spindle at stimulation onset, post-spindle stimulation produced a clear EEG response (Figure 5D). At Cz, a two-sided cluster-based permutation test against zero on the stimulation-locked time-frequency representation identified four significant clusters (N = 26, 5,000 permutations; Figure 5E). This panel displays all significant clusters: warm colors mark positive t-values, and cool colors mark negative t-values. Two clusters extended across both pre-and post-stimulation periods: a broadband cluster (1-38 Hz, −1.71 to 2.41 s; p < .001) and a sigma/beta cluster (11.5-29.5 Hz, −0.48 to 2.50 s; p < .001). A third cluster was confined to the pre-stimulation period (1-31 Hz, −2.50 to −1.55 s; p < .001), consistent with activity related to the target spindle before the stimulus. A fourth cluster appeared entirely after stimulation onset in the beta range (17.5-30.5 Hz, 0.63 to 2.08 s; p = .029), indicating a stimulation-related response that could not be attributed to the preceding spindle. In the slow-oscillation band (0.5-4 Hz, 0.5-2.0 s), post-stimulation power increased broadly across the scalp (Figure 5C): 50 of 55 channels reached significance after FDR correction (q < .05), with the strongest effects over frontal sites (AF3: z = 4.13, AF4: z = 4.40, F4: z = 4.25). Thus, post-spindle stimulation produced widespread cortical activity during the refractory period without generating a new spindle.

### 3.5. Spindle density predicts motor memory consolidation

We next asked whether participants with higher spindle density showed larger motor gains in both stimulation and adaptation naps. This question follows from prior work linking sleep spindles to motor memory consolidation (Barakat et al., 2011; Morin et al., 2008). We first tested the spindle-memory relationship within each nap using Pearson correlations, then used a linear mixed model to test whether this relationship changed between the adaptation and stimulation sessions.

During the adaptation nap session (Day 1), spindle density was positively correlated with FTT accuracy change (r = 0.508, 95% CI [0.141, 0.752], permutation p = .010, N = 25; Figure 5F). Thus, participants with higher spindle density showed stronger motor memory consolidation when no stimulation was delivered. This relationship was not present in the stimulation session: same-session Day 2 spindle density did not predict Day 2 FTT change (r = 0.013, 95% CI [-0.376, 0.399], permutation p = .948, N = 26), and Day 1 spindle density also did not significantly predict Day 2 FTT change (r = 0.204, 95% CI [-0.208, 0.554], permutation p = .323, N = 25). A linear mixed model with a density × session interaction confirmed that the spindle-density slope differed by session: the slope was positive on Day 1 (β = 0.066, z = 3.41, p < .001) but near zero on Day 2 (density × session interaction β = −0.060, z = −2.08, p = .038). The main effect of session remained positive (β = 0.056, z = 2.16, p = .031). These results indicate that motor improvement in the stimulation session depended less on endogenous spindle density than motor improvement in the adaptation session. Consistent with this interpretation, adaptation-session spindle density was negatively correlated with the stimulation benefit, defined as Day 2 - Day 1 FTT accuracy change (r = −0.425, 95% CI [-0.702, −0.036], permutation p = .039, N = 25; Figure 5G). Participants with lower baseline spindle density showed larger Day 2 gains, whereas participants with higher baseline density gained little from stimulation.

Together, these findings suggest that endogenous spindle density during the adaptation nap tracked spontaneous motor consolidation, whereas post-spindle stimulation provided the largest gains when endogenous spindle activity was relatively low.

## Discussion

Our findings show that the sleeping brain’s response to acoustic input depends on when that input falls in the spindle cycle. Stimulation during an ongoing spindle prolonged the spindle, whereas stimulation shortly after spindle offset delayed the next spindle. This dissociation held across four sound types, indicating that timing, rather than acoustic content, accounted for the effects on spindle dynamics. Post-spindle stimulation also reduced the post-nap loss of motor sequence accuracy, with no detectable effect on word-pair recall. The motor benefit was larger in participants with lower baseline spindle density, suggesting that stimulation may provide the most support when spontaneous spindle activity is weak.

During-spindle stimulation prolonged the targeted spindle without increasing its amplitude, indicating a duration-specific response. Lustenberger et al. (2016) showed that tACS at spindle frequency increased subsequent sigma power and improved motor memory, but that protocol could not determine how individual spindles changed in real time. Acoustic closed-loop work has since shown that the effect depends on when the pulse falls within the spindle cycle (Jourde et al., 2025): pulses near spindle onset can truncate the oscillation, whereas pulses during the waning phase can leave the detected spindle largely intact and elicit later sigma activity. Our protocol differed in two ways: stimulation was delivered at the fifth spindle peak rather than at detection onset, and it lasted 2 s rather than a brief pulse. The resulting prolongation identifies a condition in which sustained input, delivered after the spindle is already established, can extend the oscillation beyond its natural duration.

Our results also clarify earlier work by Choi et al. (2019), who found that closed-loop pulses after spindle detection can improve procedural memory and evoke slow activity. In that study, pulses arrived 153 ± 300 ms after the spindle dropped below threshold, with about 20% landing during ongoing spindles and 80% after offset. This timing spread makes two responses hard to separate: stimulation that reaches an ongoing spindle may prolong the current oscillation, whereas stimulation after offset may delay the next spindle. Their reported 0.7-1.2 s spindle-band rebound could therefore combine carryover from prolonged spindles with a delayed return of spindle activity after an evoked down-state. Our parallel design separates these possibilities. In Experiment 1, stimulation at the fifth spindle peak prolonged target spindles by 267 ms relative to within-subject non-target spindles, whereas stimulation 250 ms after offset did not prolong the target spindle but delayed the next one by 132 ms within subjects. In Experiment 2, post-spindle stimulation evoked cortical slow-oscillation activity without generating a new spindle and reduced the post-nap loss of motor sequence accuracy. Thus, the present findings preserve the link between spindle-timed sound and procedural memory while separating two timing-dependent outcomes that were combined in the earlier protocol: prolongation during a spindle and delayed re-emergence after it.

The two effects were similar across four acoustic variants, indicating that the timing dissociation did not depend on sound structure. The sample size limits sensitivity to small acoustic effects, but if frequency matching were critical, sigma-band amplitude-modulated noise should have produced stronger prolongation than arrhythmic or 30 Hz noise. No such difference emerged, whereas the timing effect was large and consistent. Ngo et al. (2019) similarly found that rhythmic and arrhythmic click trains during slow-oscillation up-states produced equivalent slow-oscillation and spindle-band responses, with no rhythm-specific entrainment of endogenous spindles. In our sample, neither sound rhythm nor frequency added an effect beyond timing relative to the spindle cycle. Our protocol also differed from Ngo’s in stimulus duration. Their 500 ms click train was shorter than a typical spindle, so it could test entrainment but not prolongation. Our 2 s burst was longer than a typical spindle, allowing sensory input to continue past the spindle’s natural end. This design allowed during-spindle stimulation to extend target spindles by 267 ms, an effect that sub-spindle-duration pulses would be unlikely to detect.

Post-spindle stimulation reduced motor accuracy loss relative to the adaptation nap, and the benefit was largest in participants with lower adaptation-nap spindle density. The density-FTT slope was clear on Day 1 but near zero on Day 2, as shown by the significant density × session interaction. The mechanism of this pattern is uncertain. Stimulation may support consolidation through processes that rely less on endogenous spindle activity, giving the most benefit to participants whose spindle density is low. Part of the inverse correlation may also reflect regression toward the mean among participants whose Day 1 performance already approached the measurable limit. Separating biological compensation from this statistical contribution will require dose-response designs, sham-controlled crossover designs, and larger samples.

Post-spindle stimulation supported motor memory despite not generating a new spindle. The stimulation-locked time-frequency analysis points to one possible neural basis: stimulation evoked cortical activity during the refractory period. This response is consistent with evidence that corticospinal excitability changes around spindles (Hassan et al., 2025), indicating that the cortex can still respond to input when new spindle generation is suppressed. A contrasting result from Antony et al. (2018) helps define the boundary of this effect. In that study, targeted memory reactivation cues delivered shortly after spindle offset failed to enhance memory, whereas cues delivered approximately 2.5 s later did. Those cues carried specific memory content and may have required spindle recruitment to consolidate that content. Our pink-noise stimuli carried no memory content and did not require a new spindle. This distinction is also consistent with sleep-based memory-editing work showing that content-bearing cues or interfering memories can alter aversive memories during sleep (Xia et al., 2024). Whether stimulation during the refractory period helps or hinders memory may therefore depend on what the input requires from the spindle system: selective memory reactivation may require a new spindle, whereas non-informative auditory stimulation may not.

Several design features constrain interpretation. First, Experiment 2 used a fixed-order design because the Day 2 closed-loop system required individualized spindle-detection parameters derived from the Day 1 adaptation nap. The behavioral effect remained after controlling for alertness, sleep duration, and baseline performance, but unmeasured order-related factors cannot be fully excluded. A future crossover design with two stimulation sessions in counterbalanced order would resolve this issue. Second, both samples were predominantly female, limiting generalizability across sex/gender groups. Replication in mixed-gender or matched samples remains an important next step. Third, statistical power to detect a small declarative-memory effect (dz = 0.2) was only ∼15% in our sample, so the null result on the word-pair task cannot determine whether post-spindle stimulation benefits declarative memory. Resolving this question will require a substantially larger sample.

In sum, the sleeping brain’s response to acoustic input depends on the oscillatory state in which the input arrives. The active spindle period and the post-spindle refractory period responded to identical sounds with different physiological outcomes. Future work should test whether this state dependence extends to other sleep rhythms with phase-dependent changes in excitability. More generally, phase-specific stimulation may reveal effects that event-locked protocols miss, consistent with recent proposals that sleep-based memory interventions should consider the timing of external input relative to ongoing neural oscillations (Xia & Hu, 2026). For older adults (Helfrich et al., 2018; Muehlroth et al., 2019) and clinical populations (Astori et al., 2013; Fernandez & Lüthi, 2020) whose spindle activity is reduced or abnormal, the post-spindle window may offer a route for sleep-based memory support that does not depend on generating new spindles.

## Supplementary Figure Legends

**Figure S1.** Spindle amplitude does not differ between stimulation timing conditions (Experiment 1). (A) Peak spindle amplitude (µV) for the During and Post groups. Violin plots with individual participant data; diamonds indicate mean ± SEM. Welch t(16.7) = −0.25, p = .806, d = 0.106, n.s. (B) Epoch-level amplitude distributions for both groups. (C) Amplitude by sound type and group. Mixed ANOVA: no main effect of group (F(1,20) = 0.08, p = .787, η²p = 0.004), sound type (F(3,60) = 0.71, p = .552, η²p = 0.034), or interaction (F(3,60) = 0.81, p = .491, η²p = 0.039). n = 11 per group. Error bars indicate mean ± SEM.

**Figure S2.** Cross-session spindle characteristics across experiments. (A-C) Experiment 1 During group: spindle density, mean amplitude, and mean duration in the adaptation and stimulation naps. (D-F) Experiment 1 Post group: the same three global spindle measures. (G-I) Experiment 2: Day 1 adaptation vs. Day 2 stimulation for the same measures. Continuous spindle detection was applied consistently across sessions. Paired plots show individual participants with group summaries; no comparison survived FDR correction across the nine primary paired tests.

**Figure S3.** Finger-tapping task behavioral outcomes with outlier annotation and learning curves (Experiment 2). (A) FTT accuracy improvement (post-nap minus pre-nap) for the adaptation session (Day 1, gray) and stimulation session (Day 2, amber). Violin plots show individual participant data; diamonds indicate mean ± SEM; thin lines connect within-subject pairs. Red crosses mark data points identified as outliers by the mean ± 3 SD criterion on the Day 2-Day 1 difference score, with subject IDs annotated. One participant (sub206) was excluded from all behavioral analyses (see Methods 2.1). The statistical bracket shows the paired t-test result; each measure was analyzed as a planned behavioral outcome, and uncorrected p-values are reported. (B) Block-by-block FTT accuracy across training (blocks 1-12) and post-nap testing (blocks 13-24) for Day 1 and Day 2. Thin lines show individual participant trajectories; thick lines with markers show group mean ± SEM (shaded). The vertical dashed line marks the nap boundary between training and testing. n = 26 after outlier exclusion.

**Figure S4.** Global spindle characteristics do not differ between adaptation (Day 1) and stimulation (Day 2) sessions (Experiment 2). (A) Spindle density (spindles/min). (B) Mean amplitude (µV). (C) Mean duration (s). (D) Duration coefficient of variation (SD/mean). (E) Amplitude coefficient of variation (SD/mean). Continuous spindle detection was applied identically to both sessions. Paired box plots show individual data points, with lines connecting Day 1-Day 2 pairs; diamonds indicate mean ± SEM. No feature differed significantly between sessions (all p ≥ .158, all |dz| ≤ 0.292). Teal = Day 1 (adaptation); rose = Day 2 (stimulation). n = 25 matched participants.

**Figure S5.** Individual sigma peak frequency distributions. (A) Experiment 1: sigma peak frequency histogram with kernel density estimate for the During (teal) and Post (rose) groups. (B) Experiment 2: sigma peak frequency histogram. (C) Comparison of sigma peak distributions across experiments. Dashed vertical lines indicate the 12 Hz and 16 Hz boundaries of the default sigma band. n = 22 for Experiment 1; n = 25 for Experiment 2.

## Acknowledgments

We thank Xinger Feng for assistance with data collection.

## Data and Code Availability

Analysis code, processed data, and figure-generation scripts are publicly available on OSF at https://osf.io/rbd5h/. The closed-loop stimulation software (OpenViBE configuration files) is available from the corresponding author upon reasonable request.

## Author Contributions

T.X. and X.J. contributed equally to this work. T.X., D.C., and X.H. designed the experiments. T.X. and X.J. developed the real-time spindle detection algorithm. X.J. and X.Z. collected the data. T.X. and X.J. performed the analyses. T.X., X.J., and X.H. wrote the manuscript. All authors reviewed and commented on the manuscript.

## Supplementary Tables

**Table S1.**
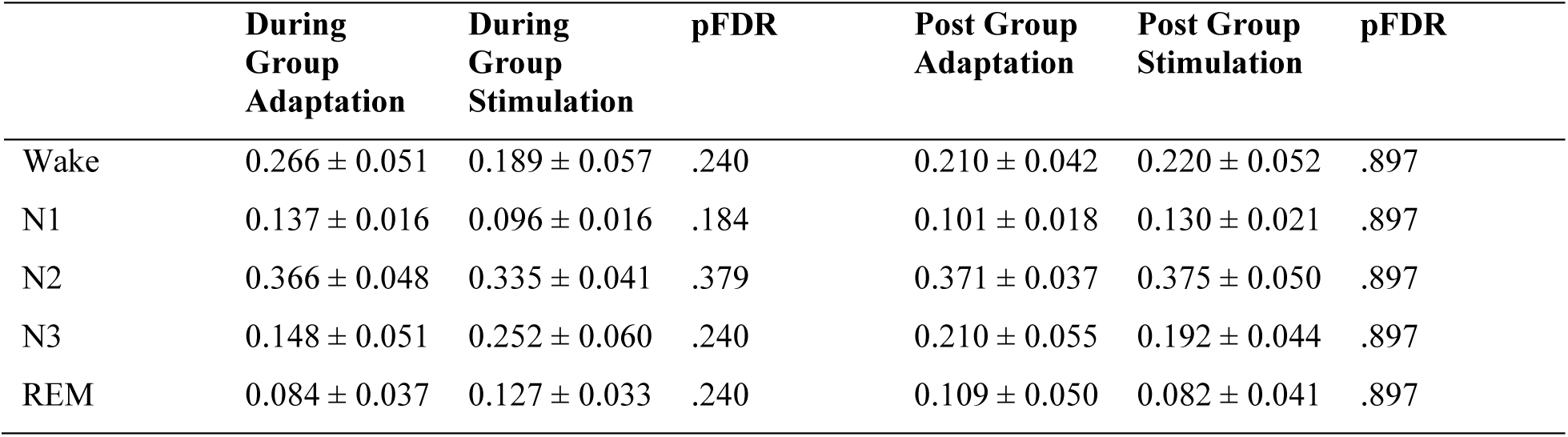
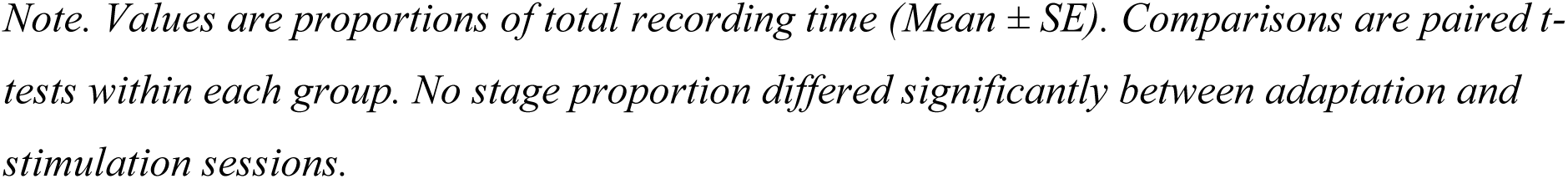
Sleep Architecture: Experiment 1 (Adaptation vs. Stimulation Nap, Paired Comparison Within Each Group)

|  | During<br>Group<br>Adaptation | During<br>Group<br>Stimulation | pFDR | Post Group<br>Adaptation | Post Group<br>Stimulation | pFDR |
| --- | --- | --- | --- | --- | --- | --- |
| Wake | 0.266 ± 0.051 | 0.189 ± 0.057 | .240 | 0.210 ± 0.042 | 0.220 ± 0.052 | .897 |
| N1 | 0.137 ± 0.016 | 0.096 ± 0.016 | .184 | 0.101 ± 0.018 | 0.130 ± 0.021 | .897 |
| N2 | 0.366 ± 0.048 | 0.335 ± 0.041 | .379 | 0.371 ± 0.037 | 0.375 ± 0.050 | .897 |
| N3 | 0.148 ± 0.051 | 0.252 ± 0.060 | .240 | 0.210 ± 0.055 | 0.192 ± 0.044 | .897 |
| REM | 0.084 ± 0.037 | 0.127 ± 0.033 | .240 | 0.109 ± 0.050 | 0.082 ± 0.041 | .897 |

*Note. Values are proportions of total recording time (Mean ± SE). Comparisons are paired *t*-tests within each group. No stage proportion differed significantly between adaptation and stimulation sessions.*

**Table S2.**
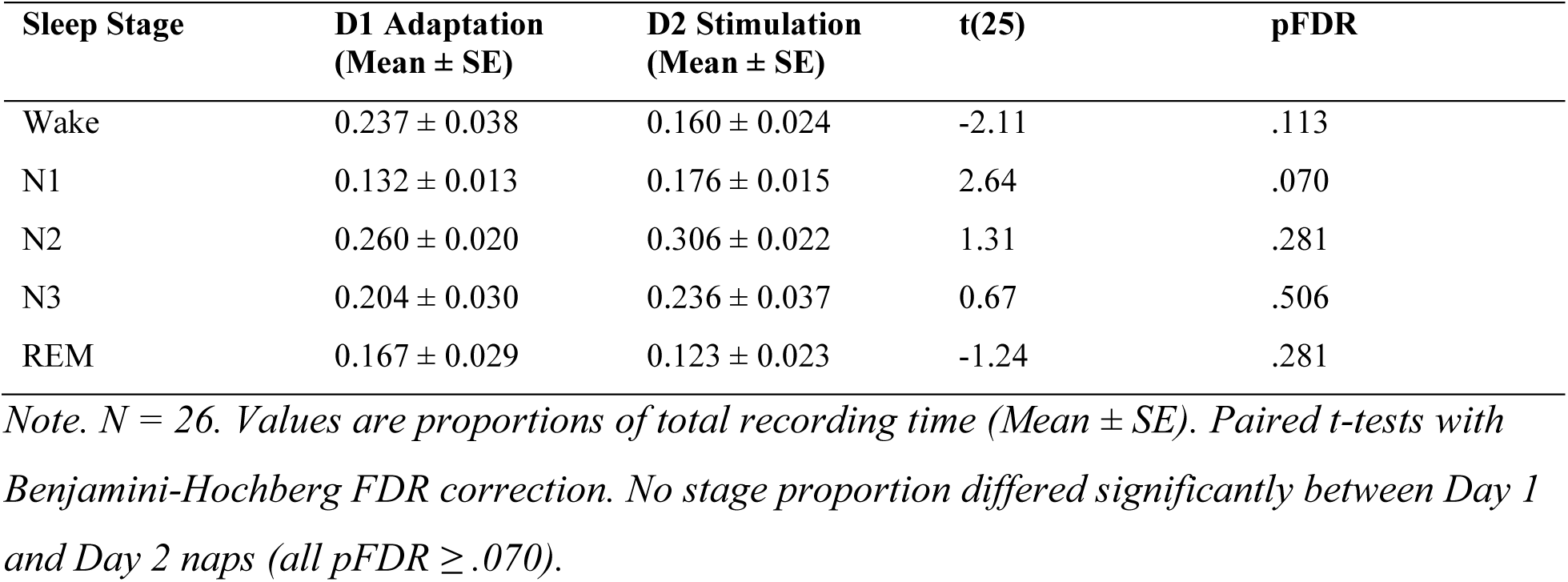
Sleep Architecture: Experiment 2 (Day 1 Adaptation vs. Day 2 Stimulation Nap)

| Sleep Stage | D1 Adaptation<br>(Mean ± SE) | D2 Stimulation<br>(Mean ± SE) | t(25) | pFDR |
| --- | --- | --- | --- | --- |
| Wake | 0.237 ± 0.038 | 0.160 ± 0.024 | -2.11 | .113 |
| N1 | 0.132 ± 0.013 | 0.176 ± 0.015 | 2.64 | .070 |
| N2 | 0.260 ± 0.020 | 0.306 ± 0.022 | 1.31 | .281 |
| N3 | 0.204 ± 0.030 | 0.236 ± 0.037 | 0.67 | .506 |
| REM | 0.167 ± 0.029 | 0.123 ± 0.023 | -1.24 | .281 |

*Note. N = 26. Values are proportions of total recording time (Mean ± SE). Paired *t*-tests with Benjamini-Hochberg FDR correction. No stage proportion differed significantly between Day 1 and Day 2 naps (all pFDR ≥ .070).*

**Table S3.** Sensitivity Analysis: Core Exp1 Results Across Spindle Amplitude Thresholds.

| Amplitude filter | N spindles | Duration During M | Duration Post M | Duration p | Latency During M | Latency Post M | Latency p |
| --- | --- | --- | --- | --- | --- | --- | --- |
| All detected (default) | 3981 | 0.925 | 0.851 | .006 | 1.069 | 1.630 | <.001 |
| Top 80% amplitude | 3185 | 0.967 | 0.881 | .005 | 1.011 | 1.593 | <.001 |
| Top 60% amplitude | 2389 | 1.001 | 0.924 | .034 | 1.134 | 1.551 | .038 |
| Top 40% amplitude | 1593 | 1.046 | 0.952 | .089 | 0.965 | 1.759 | .006 |

*Note. Core group comparisons (Welch *t*-test) were recomputed after progressively excluding low-amplitude spindles. Both the duration difference (During > Post) and the post-stimulation spindle latency difference (Post > During) remained significant across the top 60-100% of detected spindles. Loss of significance at the top 40% threshold reflects reduced statistical power (n = 11 per group) rather than a reversal of effect direction. Duration and latency values are in seconds.*

