## Supplementary figures and images for "Closed-loop auditory stimulation during sleep shapes cortical responses and memory consolidation depending on spindle timing"

### FigureS1

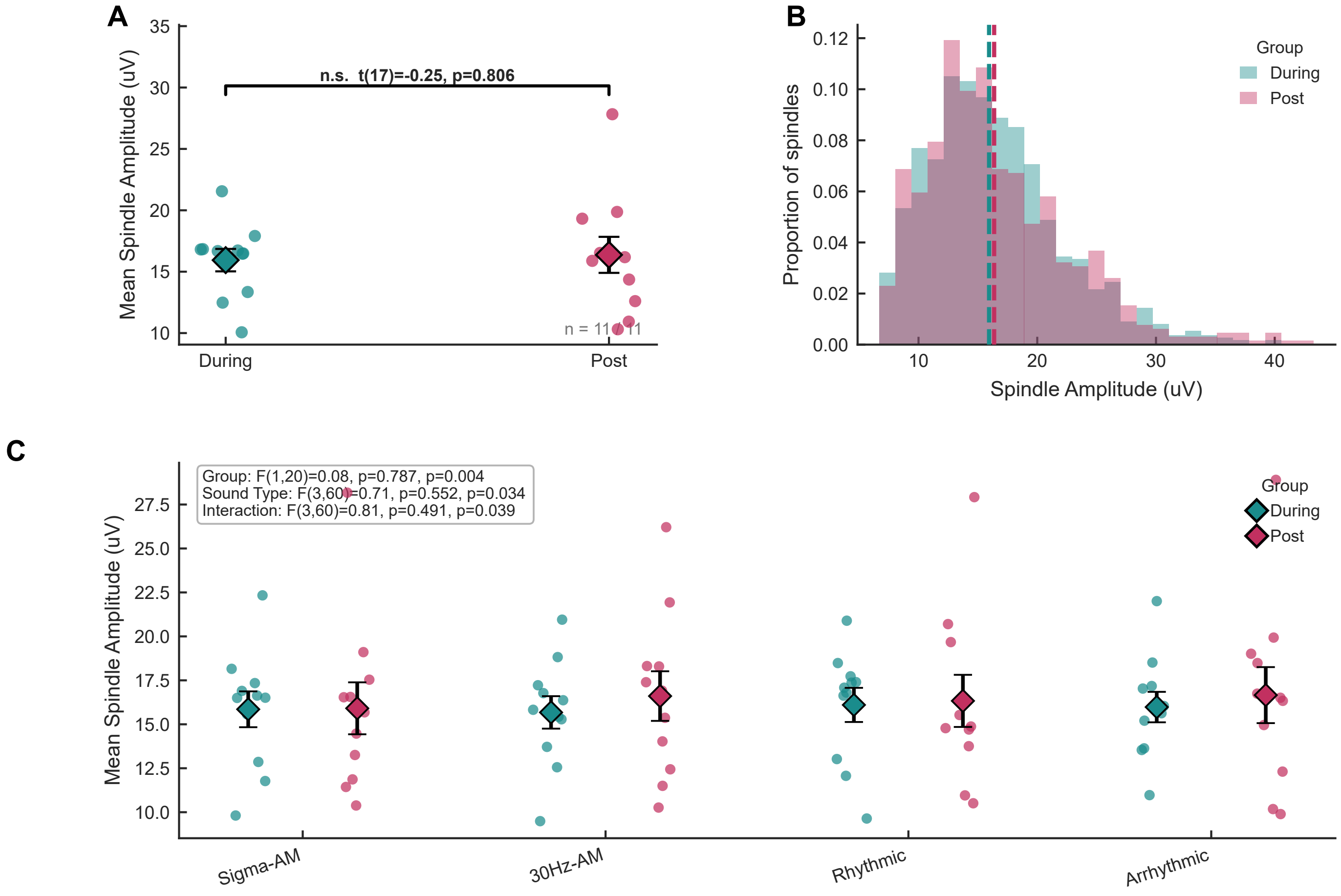

### FigureS2

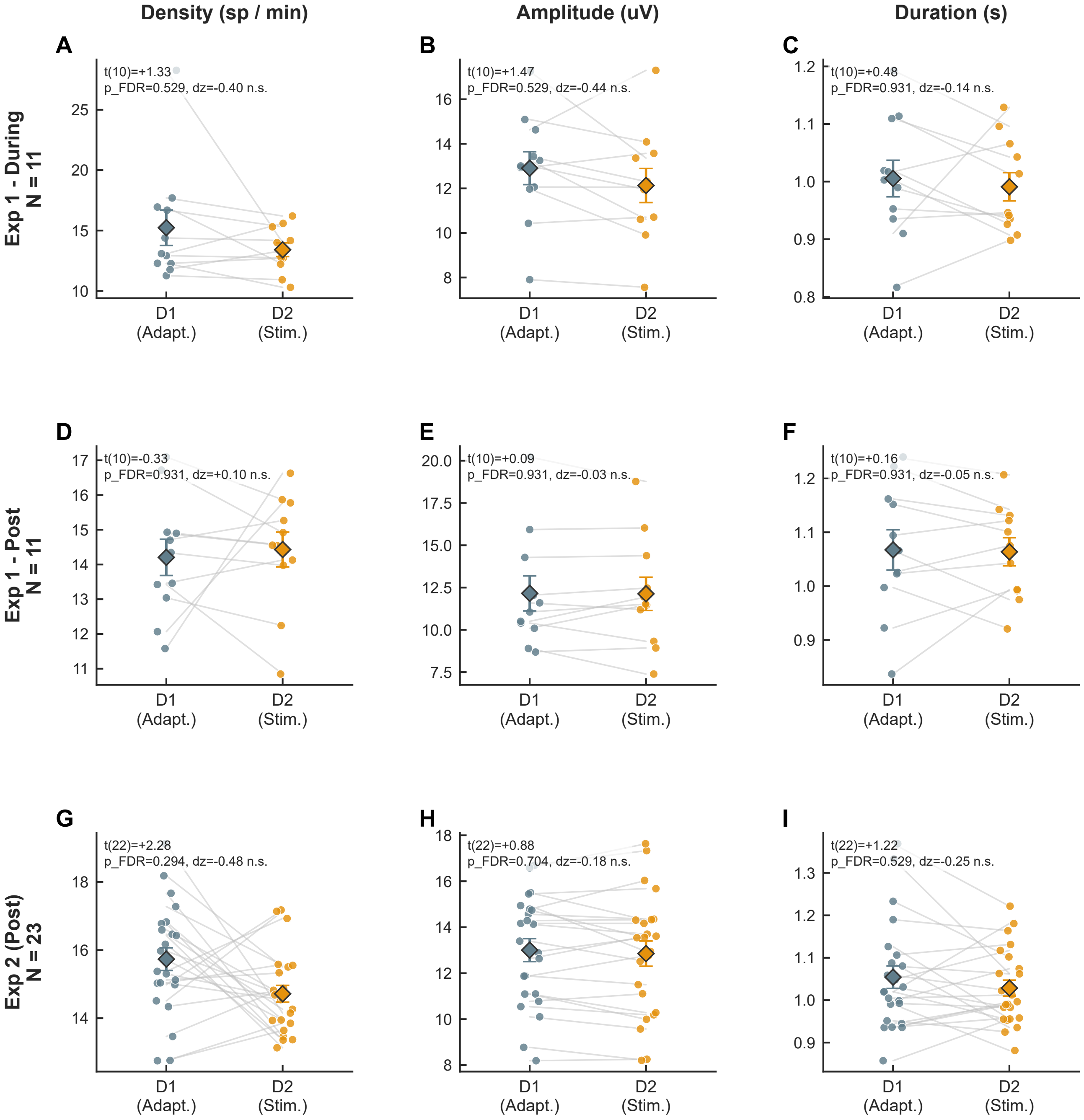

### FigureS3

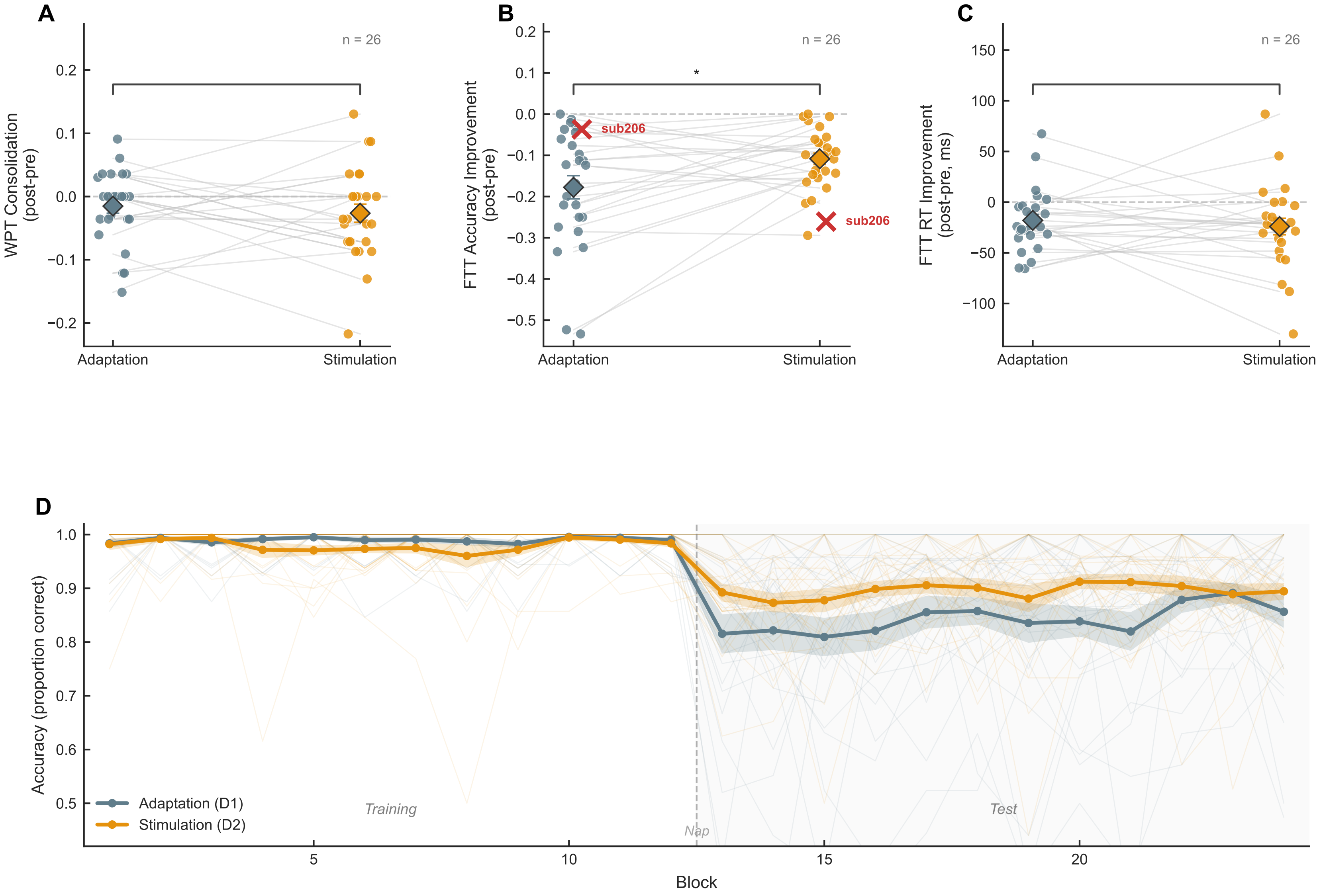

### FigureS4

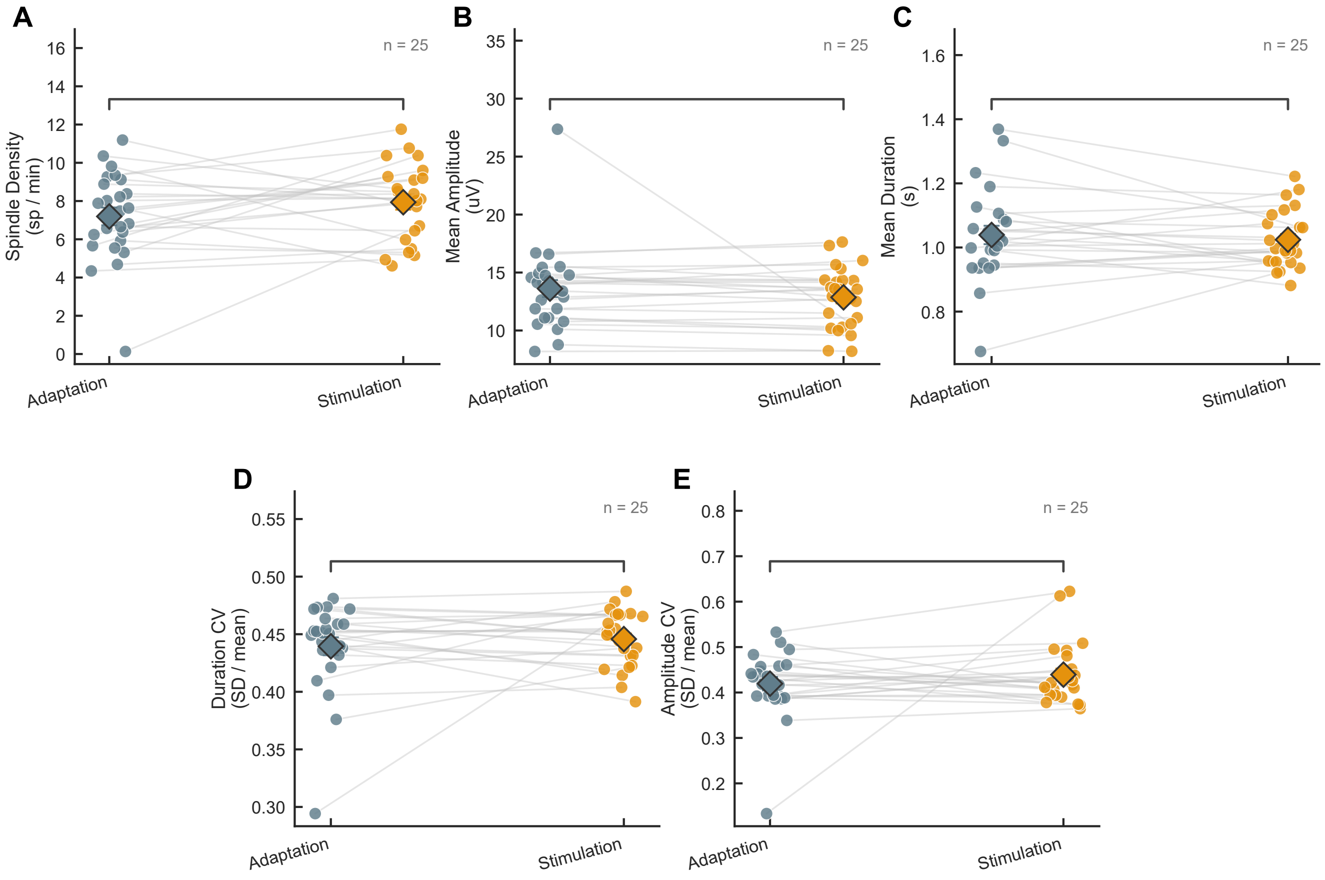

### FigureS5

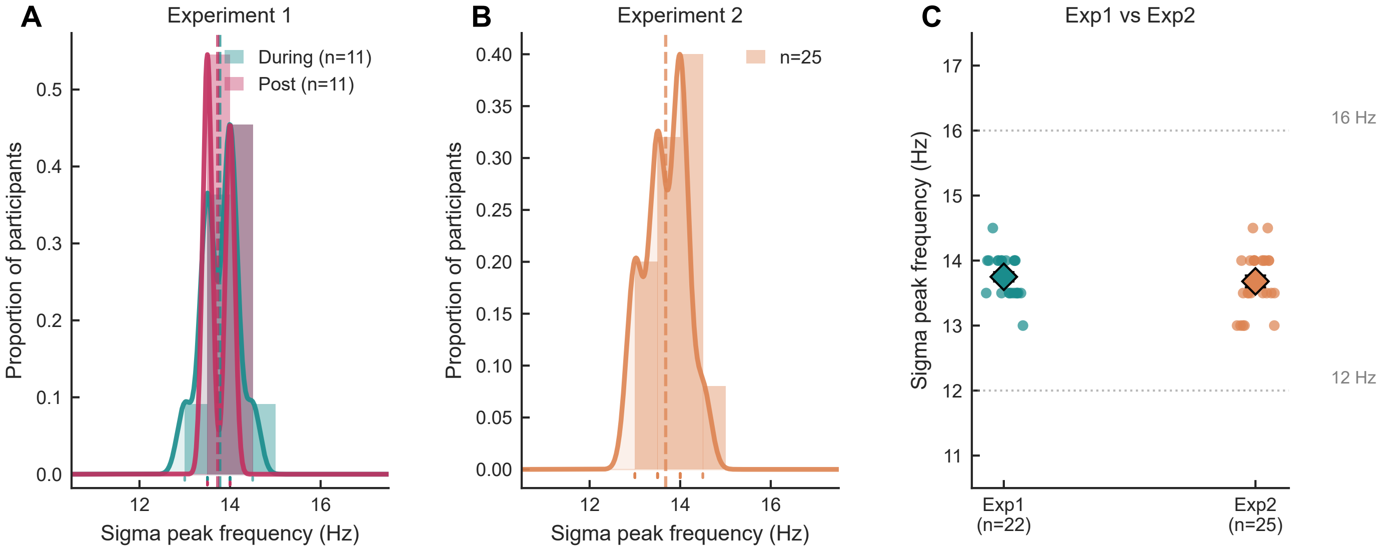
